# Inhibitory IL-10-producing CD4^+^ T cells develop in a T-bet-dependent manner and facilitate cytomegalovirus persistence via coexpression of arginase-1

**DOI:** 10.1101/2022.01.25.477742

**Authors:** Mathew Clement, Kristin Ladell, Kelly L. Miners, Morgan Marsden, Lucy Chapman, Anna Cardus Figueras, Jake Scott, Robert Andrews, Simon Clare, Valeriia V. Kriukova, Ksenia R. Lupyr, Olga V. Britanova, David R. Withers, Simon A. Jones, Dmitriy M. Chudakov, David A. Price, Ian R. Humphreys

## Abstract

Inhibitory CD4^+^ T cells have been linked with suboptimal immune responses against cancer and pathogen chronicity, but the mechanisms that underpin the development of such regulatory networks *in vivo* have remained obscure. To address this knowledge gap, we performed a comprehensive functional, phenotypic, and transcriptomic analysis of IL-10-producing CD4^+^ T cells induced by chronic infection with murine cytomegalovirus (MCMV). We identified these cells as clonally expanded and highly differentiated T_H_1-like cells that developed at sites of viral persistence in a T-bet-dependent manner and coexpressed arginase-1 (Arg1), which promotes the catalytic breakdown of L-arginine. Mice lacking Arg1-expressing CD4^+^ T cells exhibited more robust antiviral immunity and were better able to control MCMV. Conditional deletion of T-bet in the CD4^+^ lineage suppressed the development of these inhibitory cells and also enabled better immune control of MCMV. Collectively, these data elucidated the ontogeny of IL-10-producing CD4^+^ T cells and revealed a previously unappreciated mechanism of immune regulation, whereby viral persistence was facilitated by the coexpression of Arg1.

## INTRODUCTION

Immune dysregulation occurs during many persistent viral infections. High levels of ongoing viral replication, which characterize human immunodeficiency virus (HIV), hepatitis B virus (HBV), and, in mice, lymphocytic choriomeningitis virus (LCMV), typically lead to T cell exhaustion, defined by impaired effector functions, the expression of inhibitory cytokines and receptors (Wherry 2011), and substantial alterations in cellular gene expression (Doering et al. 2012). Moreover, inducible and naturally occurring FoxP3^+^ regulatory T cells accumulate during many chronic viral infections, presumably to limit immune pathology (Veiga-Parga, Sehrawat, and Rouse 2013), and T helper (T_H_)1-like cells that express the immunosuppressive cytokine IL-10 can be induced by LCMV (Parish et al. 2014), HIV (Graziosi et al. 1994), human cytomegalovirus (HCMV), and murine cytomegalovirus (MCMV) (Jones et al. 2010; Clement et al. 2016). Evidence from parasitic infections suggests that IL-10-producing T_H_1-like cells protect against immune pathology, akin to classical FoxP3^+^ regulatory T cells (Jankovic et al. 2007; Anderson et al. 2007). However, experimental deletion of IL-10 production in T cells has been shown to promote the clearance of LCMV without any obvious collateral effects (Richter et al. 2013; Parish et al. 2014; Clement et al. 2016), suggesting a potential therapeutic role for similar manipulations in humans, albeit with the possibility of an attendant risk to the development of CD8^+^ T cell memory (Laidlaw et al. 2015).

The mechanisms that induce IL-10 expression in T cells have remained unclear, despite proposed roles for costimulatory receptors, cytokines, transcription factors, and signaling via the T cell receptor (TCR) (Saraiva and O’Garra 2010). For example, chronic antigen exposure and the transcription factor Blimp-1 appear to be important for the development of IL-10-producing CD4^+^ T cells in mice infected with LCMV (Parish et al. 2014), and the inhibitory receptor TIGIT is known to act upstream of IL-10 (Schorer et al. 2020). However, it is clear that viral persistence at various tissue sites, exemplified by the salivary glands (SGs) in the context of infection with MCMV, is facilitated by IL-10 (Mandaric et al. 2012; Humphreys et al. 2007).

IFN-γ-expressing CD4^+^ T cells have been shown to limit viral replication in the SGs of mice infected with MCMV (Walton et al. 2011; Lucin et al. 1992; Jonjic et al. 1989). Nonetheless, CD4^+^ T cells also represent an important source of IL-10 (Humphreys et al. 2007; Clement et al. 2016), the production of which is promoted by IL-27 during acute infection with MCMV. In contrast, less is known about the mucosal IL-10-producing CD4^+^ T cells that appear during chronic infection with MCMV, which appear to develop independently of IL-27 (Clement et al. 2016). These cells express high levels of various transcription factors, such as c-Maf and T-bet (Clement et al. 2016), and often coexpress other molecules with putative inhibitory functions, such as Tim-3, c-Maf, Blimp-1, and IL-21 (Chihara et al. 2018; Zhu et al. 2015; Apetoh et al. 2010; Awasthi et al. 2007; Pot et al. 2009). However, the functional relevance of these characteristics has remained obscure, along with the ontogeny of IL-10-producing CD4^+^ T cells during chronic infection with MCMV.

To address these issues, we performed a comprehensive functional, phenotypic, and transcriptional analysis of IL-10-producing CD4^+^ T cells isolated from the SGs of mice infected with MCMV. Our data revealed that these cells were clonally expanded and highly differentiated T_H_1-like cells with gene expression signatures that indicated a key developmental role for T-bet. In addition, we identified an inhibitory effect attributable to arginase-1, which was coexpressed by IL-10-producing CD4^+^ T cells and facilitated the site-specific persistence of MCMV.

## RESULTS

### IL-10-producing CD4^+^ T cells display a T_H_1-like profile

To better understand the development and functionality of inhibitory CD4^+^ T cells that develop during virus chronicity, we infected with MCMV IL-10 reporter (10BiT) mice with MCMV. These mice express Thy1.1 under the IL-10 promoter (Maynard et al. 2007). At day 14 post-infection (p.i.), approximately 10–30% of CD4^+^ T cells in the SGs were Thy1.1^+^, of which ∼95% displayed an effector memory phenotype (CD44^hi^ CD62L^lo^) (**Extended Data Fig. 1a**).

We then compared the transcriptional profiles of these endogenously generated IL-10^+^ and IL-10^−^ CD4^+^ T cells, isolated via fluorescence-activated cell sorting (FACS) as Thy1.1^+^ (IL-10^+^) and Thy1.1^−^ (IL-10^−^) CD44^hi^ CD62L^lo^ CD4^+^ T cells (**Extended Data Fig. 1a**). Principal component analysis (PCA) of the RNA-seq data revealed that Thy1.1^+^ CD4^+^ T cells were transcriptionally distinct from Thy1.1^−^ CD4^+^ T cells (**Extended Data Fig. 1b**). As expected, *Il10* was highly upregulated in Thy1.1^+^ CD4^+^ T cells (**Fig. 1a**), and chromatin was more open in the *Il10* promoter region compared with Thy1.1^−^ CD4^+^ T cells (**Fig. 1b**). Genes associated with localization and cell migration (*Ccl7, Cxcl2, Cxcl12, Ccl5, Cxcl14, Ccl28, Ccl12, Ccr1, Ccr5*), cell signaling (*Ceacam1, Havcr2, Tigit, Lag3, Cd40, Cd36, Itgb4*), regulation of cellular processes (*Prdm1, Gata2, Yes1, Card10, Il33*), metabolism (*Elovl7, Galnt3, Car13, Aldh1l1, Ildrl*), including glycolysis and the tricarboxylic acid cycle (*Fbp2 and Sdhc*), oxidative phosphorylation (*osgin1*), and the mitochondrial respiratory chain (*Mt-Nd1, Ndufs6, Ndufb8, Uqcrfs1 and Uqcr11*) were also upregulated in Thy1.1^+^ CD4^+^ T cells, alongside genes associated with antiviral effector functions (*Gzmb, Prf1, Gzmk, Lyz2*) and activation (*Fgl2, Cxcr2, Nfil3*) (**Fig. 1a,c,d** and **Extended Data Fig. 1c**).

**Figure 1.**
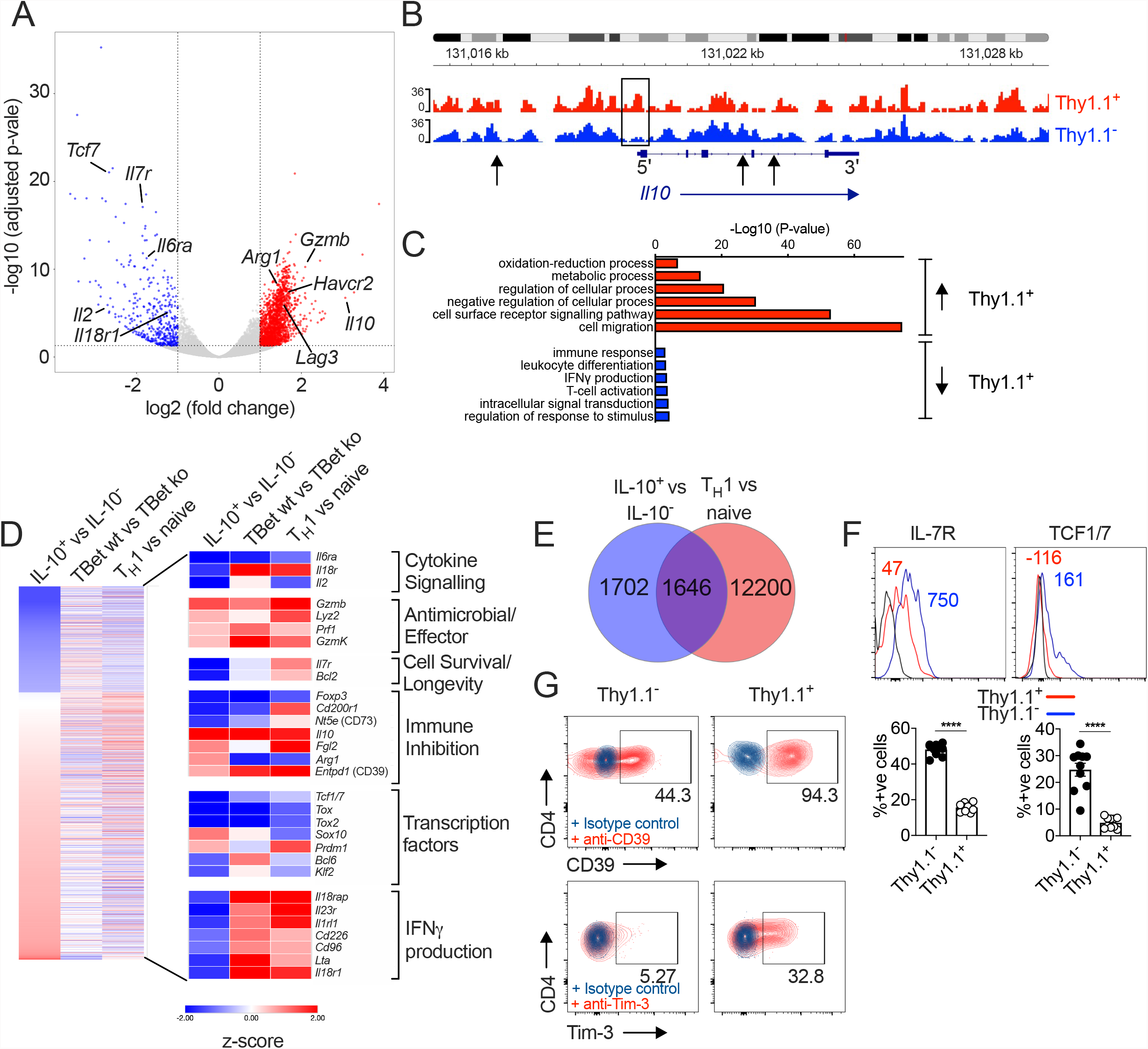
IL-10-producing CD4^+^ T cells display a T_H_1-like profile. 10BiT mice were infected with 3 × 10^4^ pfu of MCMV. Leukocytes were isolated from the SGs on day 14 p.i. and sorted as CD4^+^ CD44^+^ CD62L^−^ CD90/90.1^+^ (Thy1.1^+^) or CD90/90.1^−^ (Thy1.1^−^) populations via FACS. (**a**) Volcano plot highlighting differentially upregulated genes in Thy1.1^+^ CD4^+^ T cells (red) versus Thy1.1^−^ CD4^+^ T cells (blue). (**b**) ATAC-seq profiles showing accessible chromatin regions in the *Il10* gene for Thy1.1^+^ CD4^+^ T cells (red) and Thy1.1^−^ CD4^+^ T cells (blue). Data are shown as normalized values accounting for the total number of reads per lane. The black box indicates a major difference chromatin accessibility. Black arrows indicate binding motifs for Tbx21. (**c**) Gene ontology analysis of data from (**a**) indicating the top six modules that were upregulated (red) or downregulated (blue) in Thy1.1^+^ CD4^+^ T cells. (**d**) Heatmap comparing data from (**a**) (left column) with published data from T-bet^+^ versus T-bet-knockout CD4^+^ T cells (middle column, GSE38808) and T_H_1 versus naive CD4^+^ T cells (right column, E-MTAB-2582). (**e**) Venn diagram showing the overlap between genes enriched in Thy1.1^+^ CD4^+^ T cells (**a,d**) and genes enriched in T_H_1-like CD4^+^ T cells (E-MTAB-2582). Data in (**a**–**e**) are shown as pooled analyses from a minimum of *n* = 5 mice per group representing three independent experiments. (**f**) Representative histograms (top) and summary bar graphs (bottom) showing the expression of IL-7R and TCF1/7 among Thy1.1^+^ CD4^+^ T cells (red) and Thy1.1^−^ CD4^+^ T cells (blue). The fluorescence-minus-one control is shown in black (top). Bottom: data are shown as mean ± SEM (*n* = 10 mice per group representing two independent experiments). (**g**) Representative flow cytometry plots showing the expression of CD39 and Tim-3 among Thy1.1^+^ CD4^+^ T cells (red) versus Thy1.1^−^ CD4^+^ T cells (blue). Data are shown as pooled analyses from a minimum of *n* = 10 mice per group representing two independent experiments. \*\*\*\**P* < 0.0001 (Mann-Whitney U test).

Thy1.1^+^ CD4^+^ T cells are known to express the T_H_1-associated chemokine receptors CXCR3 and CCR5 (Clement et al. 2016). We found that MCMV-induced Thy1.1^+^ CD4^+^ T cells shared many transcripts with CD4^+^ T_H_1 cells generated *in vitro* (Stubbington et al. 2015) (**Fig. 1e**). However, genes associated with the induction of IFN-γ, including *Il18r1* and *il18rap*, were downregulated in Thy1.1^+^ CD4^+^ T cells (**Fig. 1a,c,d**), consistent with earlier functional studies of CD4^+^ T cells specific for HCMV or MCMV (Clement et al. 2016; Mason et al. 2013). Other genes that were downregulated in Thy1.1^+^ CD4^+^ T cells included *Il7* and *Tcf1/7* (**Fig. 1a,d**), which extended to the protein level (**Fig. 1f**). In addition, Thy1.1^+^ CD4^+^ T cells expressed *Entpd1* (CD39) and *Havcr2* (Tim-3) mRNA (**Fig. 1a,d**) and protein (**Fig. 1g**).

Collectively, these data showed that IL-10-producing CD4^+^ T cells exhibited a highly differentiated T_H_1-like profile during chronic infection with MCMV.

### IL-10-producing CD4^+^ T cells exhibit prominent clonal structures

IL-10-producing CD4^+^ T cells recognize a broad range of viral antigens during chronic infection with MCMV (Clement et al. 2016). To characterize these interactions in more detail and evaluate the clonal relationship between IL-10^+^ (Thy1.1^+^) and IL-10^−^ (Thy1.1^−^) CD4^+^ T cells, we used a next-generation approach to sequence the corresponding TCRs.

The repertoires of Thy1.1^+^ CD4^+^ T cells were less diverse and incorporated more prominent clonal expansions compared with the repertoires of Thy1.1^−^ CD4^+^ T cells (**Fig. 2a,b**). Several features also indicated that these expansions represented antigen-focused responses confined largely to Thy1.1^+^ CD4^+^ T cells (**Fig. 2c**–**g**). First, the number of nucleotides variants that encoded each complementarity-determining region (CDR)3α and CDR3β amino acid sequence, an indicator of antigen-specific convergence (Logunova et al. 2020), was higher overall in Thy1.1^+^ CD4^+^ T cells versus Thy1.1^−^ CD4^+^ T cells (**Fig. 2c**). Second, there were some differences in *TRBV* gene use that distinguished Thy1.1^+^ CD4^+^ T cells from Thy1.1^−^ CD4^+^ T cells, albeit with a general preference for *TRBV3, TRBV5*, and *TRBV31* (**Fig. 2d**). Third, clusters of homologous TCRβ variants, identified using a statistical model termed ALICE, (Pogorelyy et al. 2019)were detected predominantly in Thy1.1^+^ CD4^+^ T cells (**Fig. 2e**– **g**). Importantly, this latter model accounts for generation probabilities, reliably separating immunologically relevant and irrelevant but highly public TCRs.

**Figure 2.**
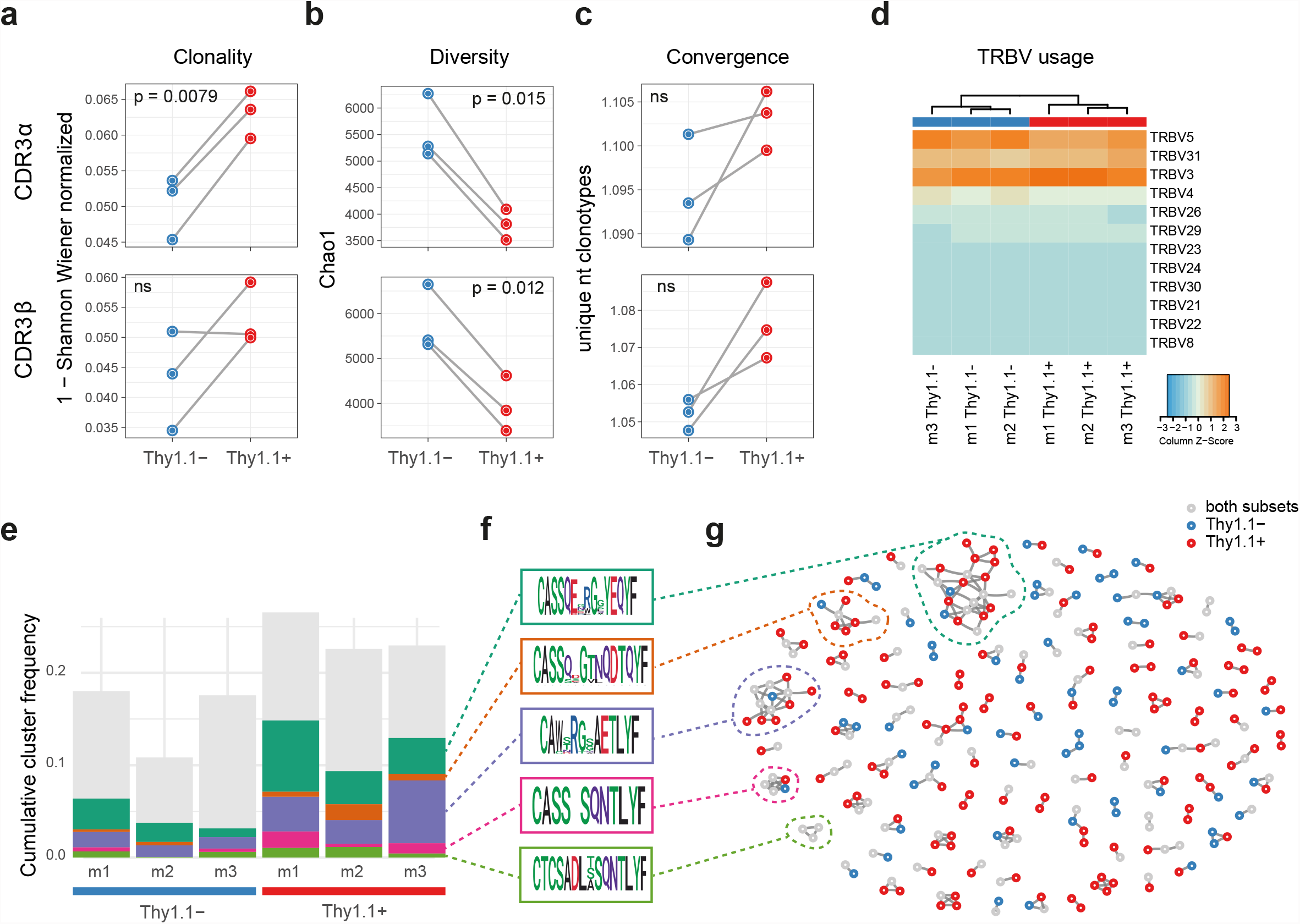
IL-10-producing CD4^+^ T cells exhibit prominent clonal structures. 10BiT mice were infected with 3 × 10^4^ pfu of MCMV. Leukocytes were isolated from the SGs on day 14 p.i. and sorted as CD4^+^ CD44^+^ CD62L^−^ CD90/90.1^+^ (Thy1.1^+^) or CD90/90.1^−^ (Thy1.1^−^) populations via FACS. (**a**) Clonality and (**b**) diversity metrics calculated for the TCRα (top) and TCRβ repertoires (bottom) derived from Thy1.1^+^ CD4^+^ T cells and Thy1.1^−^ CD4^+^ T cells. (**c**) TCR convergence measured as the average number of nucleotide sequences encoding amino acid-identical CDR3α (top) and CDR3β loops (bottom) across the 2,000 most prevalent clonotypes. (**a**–**c**) *P* values were calculated using a paired t-test with Benjamini-Hochberg correction. ns, not significant. (**d**) Hierarchical clustering of *TRBV* gene use weighted by clonotype frequency. (**e**–**g**) Cluster analysis of the 500 most prevalent TCRβ clonotypes using the tcrgarpher pipeline. (**e**) The cumulative frequency of tcrgrapher hits per sample is shown in grey. The frequency of each cluster comprising at least two tcrgrapher hits was calculated for each sample and averaged across all six repertoires. The five most prevalent clusters are shown in color. (**f**) Amino acid logos for each of five most prevalent clusters. (**g**) Visual representation of clusters comprising at least two tcrgrapher hits. Nodes represent unique amino acid sequences. Edges connect sequences with a single amino acid mismatch. Amino acid sequences present only in Thy1.1^+^ CD4^+^ T cells are shown in red, amino acid sequences present only in Thy1.1^−^ CD4^+^ T cells are shown in blue, and amino acid sequences present in both Thy1.1^+^ CD4^+^ T cells and Thy1.1^−^ CD4^+^ T cells are shown in grey. Data are shown as pooled analyses from of *n* = 4 mice per group representing three independent experiments (m1, m2, and m3).

It should be noted that none of these differences were absolute. For example, the clusters of TCRβ variants identified in Thy1.1^+^ CD4^+^ T cells also occurred at lower cumulative frequencies in Thy1.1^−^ CD4^+^ T cells (**Fig. 2e**–**g**), and the TCRα and TCRβ repertoires overlapped considerably between Thy1.1^+^ CD4^+^ T cells and Thy1.1^−^ CD4^+^ T cells (**Extended Data Fig. 2a**). Moreover, there were no prominent differences in the physicochemical properties of amino acids in the central parts of the CDR3α and CDR3β loops, which generally differ among functionally discrete subsets of CD4^+^ T cells (Kasatskaya et al. 2020), to indicate an ontogenetic divergence between Thy1.1^+^ CD4^+^ T cells and Thy1.1^−^ CD4^+^ T cells (**Extended Data Fig. 2b**).

Collectively, these data revealed the presence of common molecular signatures that predominated in Thy1.1^+^ CD4^+^ T cells, consistent with the notion of an antigen-driven process of differentiation leading to the production of IL-10.

### IL-10-producing CD4^+^ T cells promote viral persistence via coexpression of arginase-1

IL-10 expression by CD4^+^ T cells has been associated with numerous other inhibitory functions and cell membrane-expressed markers of exhaustion (Saraiva and O’Garra 2010). In line with these observations, we found that MCMV-induced Thy1.1^+^ CD4^+^ T cells expressed a module of inhibitory genes, including *Lag3, Fgl2, Havcr2*, and *Entpd1* (**Fig. 1d**). We also detected significant upregulation of *Arg1* (**Fig. 1d**). Arginase-1 (Arg1) promotes the catalytic breakdown of L-arginine (Munder 2009) and has been shown to inhibit the proliferation of T cells (Czystowska-Kuzmicz et al. 2019; Rodriguez et al. 2002; Rodriguez et al. 2004).

Arg1 expression by T cells has been reported previously (Washburn et al. 2019). The functional relevance of this observation has nonetheless remained obscure. To address this knowledge gap, we first confirmed the open chromatin structure around *Arg1* in Thy1.1^+^ CD4^+^ T cells (**Fig. 3a**). We also confirmed expression at the protein level via flow cytometry (**Fig. 3b**) and Western blotting (**Fig. 3c**) in experiments incorporating control mice lacking the ability to express Arg1 in the CD4^+^ lineage (*CD4*^cre+^*Arg1*^flox+/+^). Our data showed that Arg1 was expressed by CD4^+^ T cells almost exclusively in the SGs (**Fig. 3b,c**).

**Figure 3.**
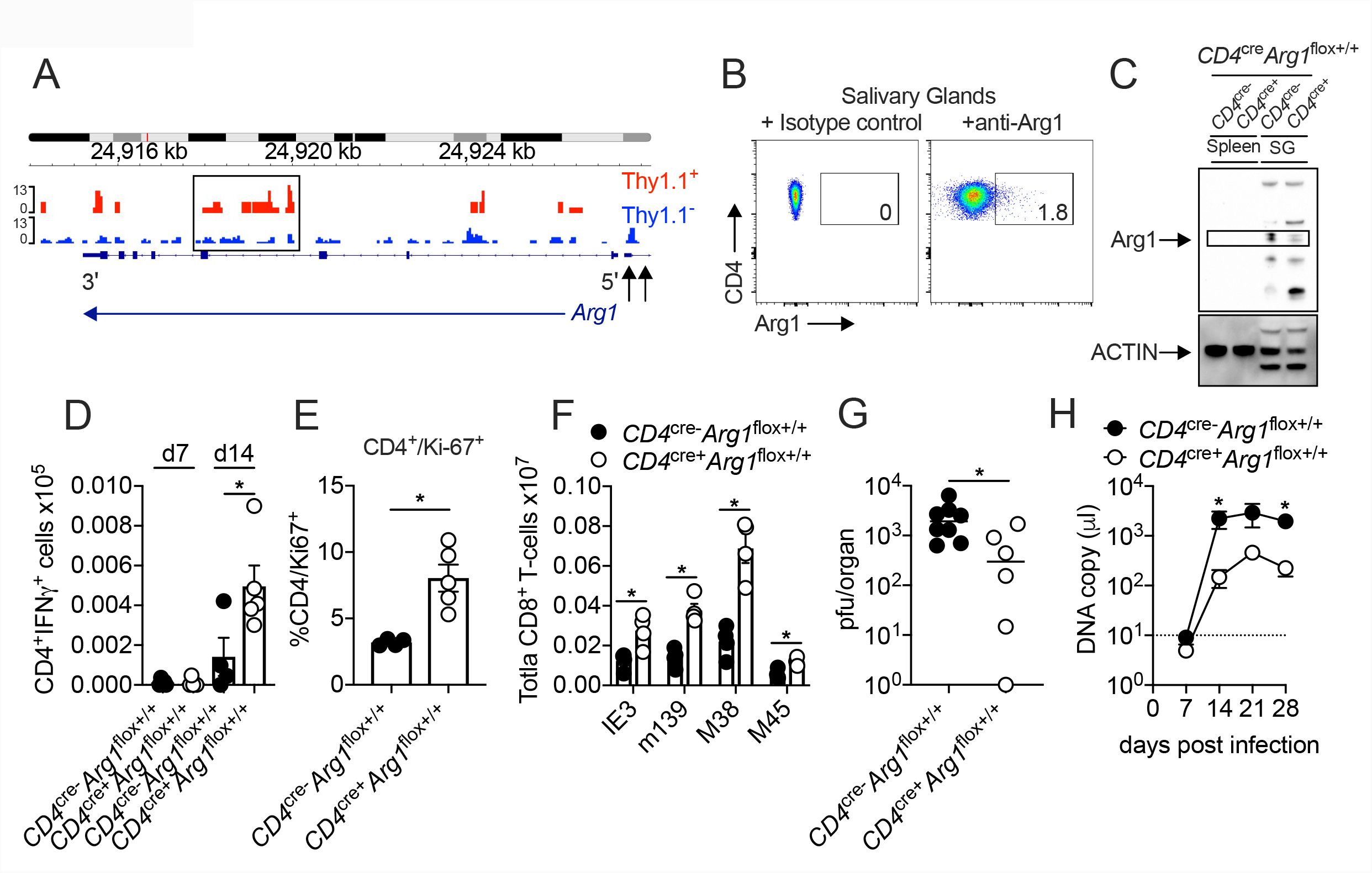
IL-10-producing CD4^+^ T cells promote viral persistence via coexpression arginase-1. (**a**) 10BiT mice were infected with 3 × 10^4^ pfu of MCMV. Leukocytes were isolated from the SGs on day 14 p.i. and sorted as CD4^+^ CD44^+^ CD62L^−^ CD90/90.1^+^ (Thy1.1^+^) or CD90/90.1^−^ (Thy1.1^−^) populations via FACS. ATAC-seq profiles showing accessible chromatin regions in the *Arg1* gene for Thy1.1^+^ CD4^+^ T cells (red) and Thy1.1^−^ CD4^+^ T cells (blue). Data are shown as normalized values accounting for the total number of reads per lane. The black box indicates a major difference in chromatin accessibility. Black arrows indicate binding motifs for Tbx21. Data are shown as pooled analyses from a minimum of *n* = 5 mice per group representing three independent experiments. (**b**) Representative flow cytometry plots showing the expression of Arg1 among CD4^+^ T cells isolated from the SGs on day 14 p.i. (**c**) Expression of Arg1 among leukocytes isolated via magnetic separation from the SGs or spleens of *CD4*^cre−^*Arg1*^flox+/+^ or *CD4*^cre+^*Arg1*^flox+/+^ mice on day 14 p.i. detected by Western blot. (**b,c**) Data are shown as pooled analyses from a minimum of *n* = 7 mice per group representing three independent experiments. (**d**) MCMV-specific CD4^+^ T cell responses in the SGs on days 7 and 14 p.i. measured using flow cytometry to detect IFN-γ. Immunodominant peptides were pooled for stimulation. Data are shown as mean ± SEM (*n* = 4–6 mice per group representing three independent experiments). (**e**) Expression of Ki-67 among CD4^+^ T cells isolated from the SGs on day 14 p.i. measured via flow cytometry. Data are shown as mean ± SEM (*n* = 4–5 mice per group representing two independent experiments). (**f**) MCMV tetramer^+^ CD8^+^ T cells quantified in spleens on day 30 p.i. via flow cytometry. Data are shown as mean ± SEM (*n* = 4 mice per group representing two independent experiments). (**g**) MCMV replication in SG homogenates on day 30 p.i. measured via plaque assay. Data are shown as individual points with median values (*n* = 6 mice per group representing two or three independent experiments). (**h**) Viral genomes in saliva on days 7, 14, 21, and 28 p.i. measured via qPCR. Data are shown as mean ± SEM (*n* = 8 mice per group representing two independent experiments). \**P* < 0.05 (Mann-Whitney U test).

We then infected *CD4*^cre−^*Arg1*^flox+/+^ and *CD4*^cre+^*Arg1*^flox+/+^ mice with MCMV. Higher numbers of IFN-γγγ-expressing CD4^+^ T cells (**Fig. 3d**) and higher frequencies of proliferating (Ki-67^+^) CD4^+^ T cells (**Fig. 3e**) were observed after viral antigen stimulation in the SGs of *CD4*^cre+^*Arg1*^flox+/+^ versus *CD4*^cre−^*Arg1*^flox+/+^ mice during the chronic phase of infection with MCMV. Similarly, higher numbers of virus-specific CD8^+^ T cells, quantified using tetrameric antigen probes, were detected in the spleens of *CD4*^cre+^*Arg1*^flox+/+^ versus *CD4*^cre−^*Arg1*^flox+/+^ mice on day 30 p.i. (**Fig. 3f**).

IFN-γγγ-expressing CD4^+^ T cells are known to restrict MCMV replication in the SGs (Walton et al. 2011). Accordingly, we found that viral replication in the SGs (**Fig. 3g**) and shedding in the saliva (**Fig. 3h**) were reduced in *CD4*^cre+^*Arg1*^flox+/+^ versus *CD4*^cre−^*Arg1*^flox+/+^ mice during the chronic phase of infection with MCMV. In contrast, deletion of *Arg1* in myeloid cells, achieved using *LysM*^cre+^*Arg1*^flox+/+^ mice, had no impact on the replication or shedding of MCMV (**Extended Data Fig. 3a,b**). Of note, there was no evidence that *CD4*^cre+^*Arg1*^flox+/+^ mice were more susceptible to virus-induced autoimmunity, as indicated by anti-Sjögrens syndrome antigen (SSA) titers comparable to those observed in *CD4*^cre−^*Arg1*^flox+/+^ mice (**Extended Data Fig. 3c**).

Collectively, these data indicated that Arg1 expression by CD4^+^ T cells selectively inhibited the accumulation of virus-specific CD4^+^ and CD8^+^ T cells during chronic infection with MCMV.

### IL-10-producing CD4^+^ T cells develop in a T-bet-dependent manner

In line with our previous work (Clement et al. 2016), we noted that Thy1.1^+^ CD4^+^ T cells expressed T-bet more commonly at the protein level than Thy1.1^−^ CD4^+^ T cells (**Fig. 4a**), and concordantly, we found that open chromatin was enriched in the *Tbx21* region of Thy1.1^+^ CD4^+^ T cells versus Thy1.1^−^ CD4^+^ T cells (**Fig. 4b**). We also detected considerable overlap between the gene expression profiles of Thy1.1^+^ CD4^+^ T cells and T-bet-orchestrated CD4^+^ T_H_1 cells generated *in vitro* (Zhu et al. 2012) (**Fig. 4c**). Moreover, T-bet suppresses the expression of TCF1/7 and IL-7R (Oestreich, Huang, and Weinmann 2011; Dominguez et al. 2015), mimicking key phenotypic features of Thy1.1^+^ CD4^+^ T cells (**Fig. 1d,f**). We therefore hypothesized that T-bet promoted the development of IL-10-producing, Arg1-expressing CD4^+^ T cells during chronic infection with MCMV.

**Figure 4.**
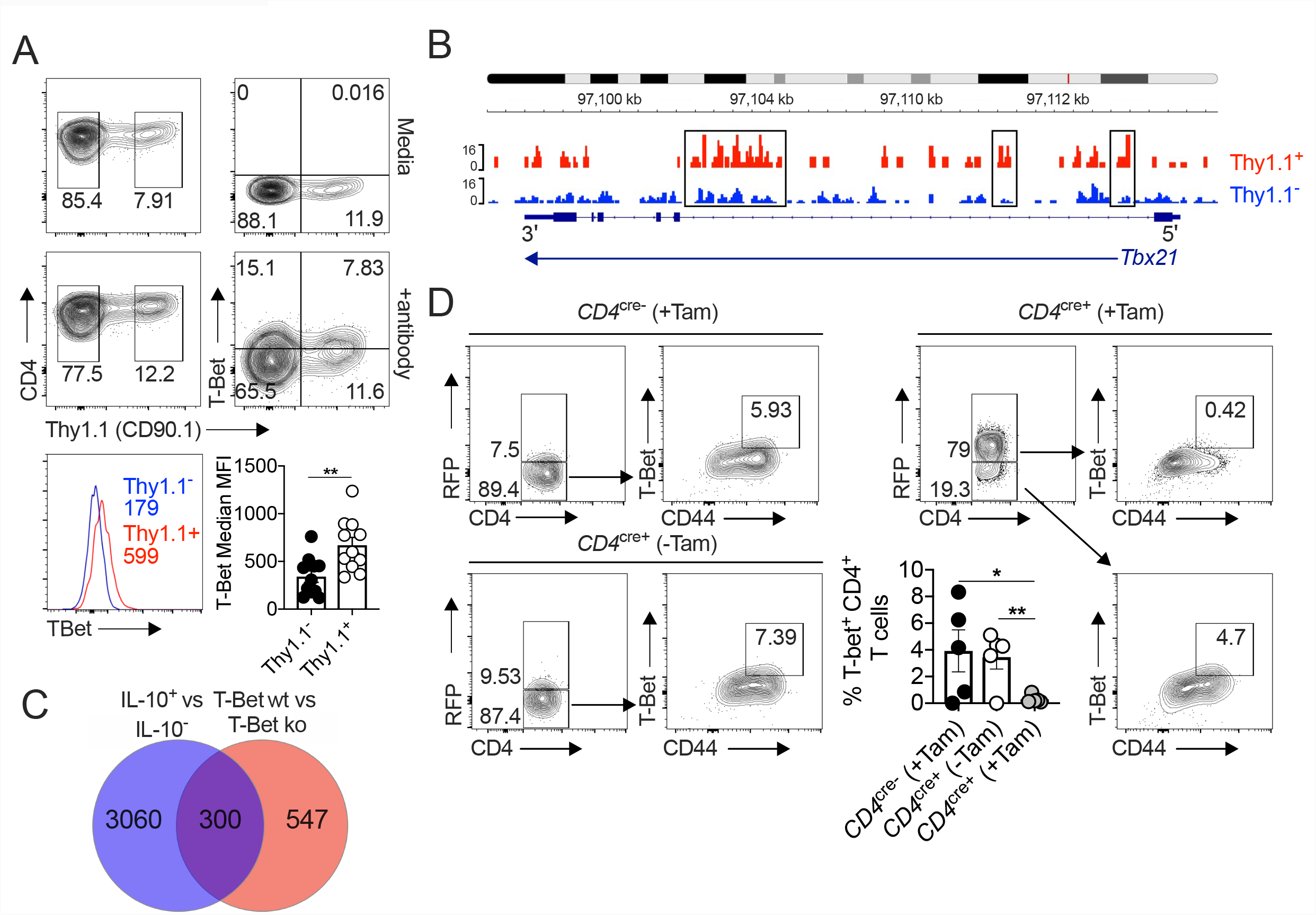
IL-10-producing CD4^+^ T cells express T-bet and T-bet-inducible genes. (**a,b**) 10BiT mice were infected with 3 × 10^4^ pfu of MCMV. (**a**) Top: representative flow cytometry plots showing the expression of T-bet versus Thy1.1 among leukocytes isolated from the SGs on day 14 p.i. Bottom: representative histograms (left) and summary bar graph (right) showing the expression of T-bet among Thy1.1^+^ CD4^+^ T cells (red) and Thy1.1^−^ CD4^+^ T cells (blue). Data are shown as mean ± SEM (*n* = 5–6 mice per group pooled from two independent experiments). MFI, median fluorescence intensity. (**b**) Leukocytes were isolated from the SGs on day 14 p.i. and sorted as CD4^+^ CD44^+^ CD62L^−^ CD90/90.1^+^ (Thy1.1^+^) or CD90/90.1^−^ (Thy1.1^−^) populations via FACS. ATAC-seq profiles show accessible chromatin regions in the *Tbx21* gene for Thy1.1^+^ CD4^+^ T cells (red) and Thy1.1^−^ CD4^+^ T cells (blue). Data are shown as normalized values accounting for the total number of reads per lane. The black boxes indicate major differences in chromatin accessibility. Data are shown as pooled analyses from a minimum of *n* = 5 mice per group representing three independent experiments. (**c**) Venn diagram showing the overlap between genes enriched in Thy1.1^+^ CD4^+^ T cells (**Fig. 1a,d**) and genes enriched in T-bet^+^ CD4^+^ T cells (GSE38808). (**d**) *CD4*^cre−^/^ERT2^-*Tbx21*^flox/++^ (*CD4*^cre−^) and *CD4*^cre+^/^ERT2^-*Tbx21*^flox/++^ (*CD4*^cre+^) mice were infected with 3 × 10^4^ pfu of MCMV. Tamoxifen was administered (+Tam) or withheld (−Tam) from days 7 to 12 p.i. Leukocytes were isolated from the SGs on day 14 p.i. Representative flow cytometry plots show the expression of CD4 versus RFP (left) and CD44 versus T-bet for the gated populations (right). Data in the bar graph are shown as mean ± SEM (*n* = 4–5 mice per group representing four or five independent experiments). \**P* < 0.05, ** *P* < 0.01 (Mann-Whitney U test).

To test this notion, we crossed tamoxifen-inducible *CD4*^cre+^ mice with *Tbx21*^flox^ mice (Intlekofer et al. 2008), allowing us to suppress T-bet expression at the onset of viral chronicity (day 7 p.i.). These mice were further crossed to incorporate a *Rosa26*^flSTOPtdRFP^ allele (Luche et al. 2007), enabling the identification of cells in which *Cre* was expressed via the detection of tandem-dimer red fluorescent protein (tdRFP). Mice were then infected with MCMV. Tamoxifen was administered daily for 5 d from day 7 p.i. to deplete T-bet in a CD4-dependent manner, clearly associating with the coincident expression of RFP (**Fig. 4d** and **Extended Data Fig. 3d**).

CD4-specific T-bet depletion reduced the accumulation of Arg1-expressing CD4^+^ T cells by day 14 p.i. (**Fig. 5a**). However, it seemed unlikely that T-bet directly stimulated the expression of IL-10 and Arg-1, because the corresponding binding motifs were not preferentially accessible in the *Il10* and *Arg1* regions (**Figs. 1b** and **3b**). Instead, we observed a shift towards a less differentiated phenotype in *CD4*^cre+^*Tbx21*^flox^ mice (**Fig. 5c–f)**, with decreased expression of CD44 (**Fig. 5c**) and PD-1 (**Fig. 5d**) and increased expression of CD62L (**Fig. 5e**) and IL-7R (**Fig. 5f**) in the absence of T-bet. In addition, T-bet depletion was associated with increased numbers of virus-specific IFN-γ-expressing CD4^+^ T cells (**Fig. 5g**) and enhanced control of viral replication in the SGs (**Fig. 5h**), as well as reduced salivary shedding of MCMV (**Fig. 5i**). In accordance with the observation that IL-10-producing CD4^+^ T cells occur transiently during the early stages of viral chronicity (Clement et al. 2016), we also found that CD4-specific T-bet depletion was no longer protective by day 28 p.i. (**Fig. 5h,i**).

**Figure 5.**
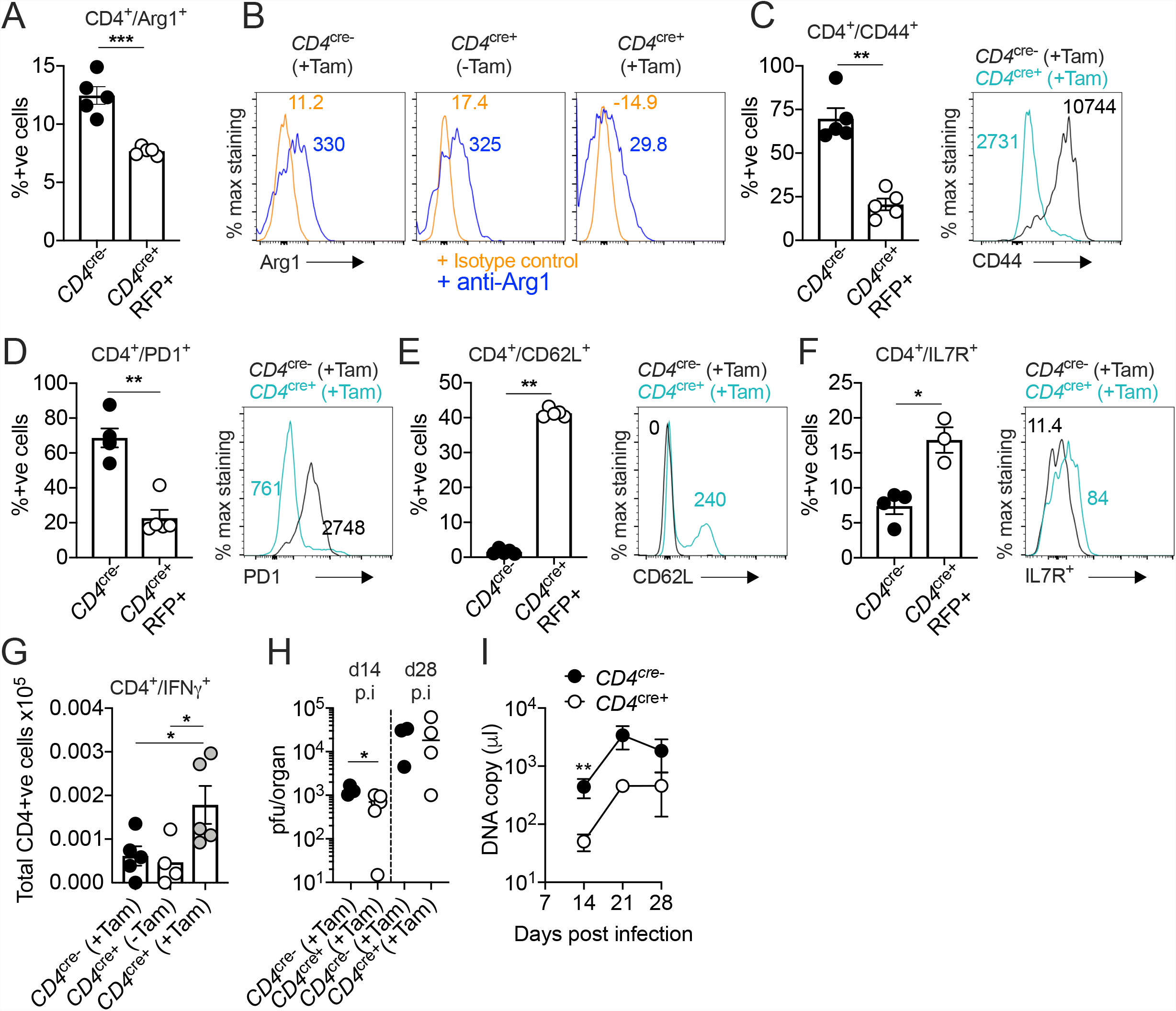
IL-10-producing CD4^+^ T cells develop in a T-bet-dependent manner. (**a**–**i**) *CD4*^cre−^/^ERT2^-*Tbx21*^flox/++^ (*CD4*^cre−^) and *CD4*^cre+^/^ERT2^-*Tbx21*^flox/++^ (*CD4*^cre+^) mice were infected with 3 × 10^4^ pfu of MCMV. Tamoxifen was administered (+Tam) or withheld (−Tam) from days 7 to 12 p.i. Leukocytes were isolated from the SGs on day 14 p.i. (**a,b**) Summary bar graph (**a**) and representative histograms (**b**) showing the expression of Arg1 among CD4^+^ T cells measured via flow cytometry. (**c**–**g**) Summary bar graphs (left) and representative histograms (right) showing the expression of CD44 (**c**), PD-1 (**d**), CD62L (**e**), and IL-7R (**f**) among CD4^+^ T cells measured via flow cytometry. Data in (**c**–**e**) are shown as mean ± SEM (*n* = 5 mice per group representing two independent experiments). Data in (**f**) are shown as mean ± SEM (*n* = 3–4 mice per group representing two independent experiments). (**g**) MCMV-specific CD4^+^ T cell responses in the SGs measured using flow cytometry to detect IFN-γ. Immunodominant peptides were pooled for stimulation. Data are shown as mean ± SEM (*n* = 4–5 mice per group representing two independent experiments). (**h**) MCMV replication in SG homogenates on days 14 and 28 p.i. measured via plaque assay. Data are shown as individual points with median values (*n* = 3–6 mice per group representing three independent experiments). (**i**) Viral genomes in saliva on days 7, 14, 21, and 28 p.i. measured via qPCR. Data are shown as mean ± SEM (*n* = 5 mice per group representing two independent experiments). Data in (**a**) and (**c**–**f**) show all groups after the administration of tamoxifen. \**P* < 0.05, \*\**P* < 0.01, \*\*\**P* < 0.001 (Mann-Whitney U test).

Collectively, these data suggested that recurrent antigen stimulation drove the observed T-bet-mediated differentiation of CD4^+^ T cells, leading to coincident expression of IL-10 and Arg1 during chronic infection with MCMV.

## DISCUSSION

In this study, we identified IL-10-producing CD4^+^ T cells as highly differentiated T_H_1-like cells, which occurred transiently at mucosal sites of viral persistence under the strict governance of T-bet. These cells also expressed Arg1. Further investigation revealed that Arg1 expression was a critical function of IL-10-producing CD4^+^ T cells, suppressing antiviral immunity and facilitating viral chronicity in the context of infection with MCMV.

Our finding that *Arg1* was one of the putative immune regulatory genes expressed by IL-10-producing CD4^+^ T cells was initially counterintuitive, given the reported lack of Arg1 protein expression by human CD4^+^ T cells *in vitro* (Geiger et al. 2016). However, *Arg1* expression by CD4^+^ and CD8^+^ T cells has been described in patients with sepsis (Washburn et al. 2019), indicating a potential role for generic infectious stimuli and/or an antigen-driven process mediated via the TCR. The latter possibility is consistent with the repertoire data, which revealed that prominent clonal expansions were associated with the production of IL-10. Importantly, we confirmed that Arg1 was expressed at the protein level, although the signals were relatively weak in flow cytometry experiments, consistent with an earlier study of patients infected with HBV (Pallett et al. 2015). It was also notable that Arg1 protein expression was confined to CD4^+^ T cells isolated from the SGs during chronic but not acute infection with MCMV.

L-arginine deficiency results in cell cycle arrest via the downregulation of CD3β, a key component of the TCR (Rodriguez et al. 2002), and T cell proliferation is impaired in the absence of L-arginine (Rodriguez, Quiceno, and Ochoa 2007). In line with these observations, we found that improved control of viral replication in mice lacking Arg1^+^ CD4^+^ T cells was associated with increased numbers of MCMV-specific effector CD4^+^ T cells, which are known to limit viral replication in the SGs (Walton et al. 2011; Lucin et al. 1992). Accordingly, the acquisition of Arg1 expression by highly differentiated CD4^+^ T cells impinges on antiviral immunity, representing an inhibitory mechanism that operates alongside the production of IL-10 (Humphreys et al. 2007; Clement et al. 2016).

IL-10 is regulated by a number of transcription factors, including those required for T cell differentiation, and is induced downstream of the TCR (Saraiva, Vieira, and O’Garra 2020). Our discovery that T-bet promotes the development of IL-10-producing CD4^+^ T cells aligns with the concept that chronic antigen stimulation can trigger a negative feedback loop via these mechanisms, which concurrently drive activation and differentiation. It nonetheless remains less clear to what extent T-bet directly orchestrates the expression of other inhibitory molecules by CD4^+^ T cells. There was no notable increase in chromatin-accessible binding motifs for T-bet within either the *Il10* or *Arg1* genes in IL-10-producing CD4^+^ T cells. However, we did find that deletion of T-bet after acute infection limited the accumulation of highly differentiated CD4^+^ T cells, including those expressing Arg1 and IL-10. In line with this observation, which suggested a differentiation-linked process of functional remodelling driven by chronic antigen exposure, T-bet is known to promote the differentiation of CD8^+^ T cells (Dominguez et al. 2015; Omilusik et al. 2015) and inhibit the expression of TCF1/7 (Omilusik et al. 2015) and IL-7R (Intlekofer et al. 2007), both of which are downregulated in IL-10^+^ Arg1^+^ CD4^+^ T cells induced by MCMV.

The mechanisms that directly induce the expression of *Il10* and *Arg1* in CD4^+^ T cells remain obscure. Although IL-10-producing CD4^+^ T cells were characterized by prominent clonal expansions, repertoire analysis revealed no obvious determinative role for the TCR. The transcription factor c-Maf, which promotes the expression of IL-10 in multiple T_H_ cell subsets (Gabrysova et al. 2018), is upregulated in IL-10-producing CD4^+^ T cells during chronic infection with MCMV (Clement et al. 2016). We also demonstrated previously that ICOS signaling promotes the accumulation of IL-10-producing CD4^+^ T cells under the same conditions (Clement et al. 2016), and c-Maf is downstream of ICOS (Bauquet et al. 2009; Nurieva et al. 2003). It therefore appears likely that an ICOS-c-Maf axis participates in the direct induction of *Il10*, although it is less clear how this applies to *Arg1*.

In summary, we have demonstrated that T-bet activity during a chronic viral infection can impede antiviral immune control by driving the development of highly differentiated CD4^+^ T cells that express genes encoding inhibitory molecules, including IL-10 and Arg1. These observations are conceivably important not only from a biological perspective but also from a translational perspective, revealing from a previously unappreciated mechanism through which CD4^+^ T cells can suppress potentially harmful immune responses via the regulation of L-arginine.

## ONLINE METHODS

### Mice

10BiT reporter mice were originally derived by Padraic Fallon (Trinity College Dublin). These mice express CD90/CD90.1 (Thy1.1) under the control of the IL-10 promoter and retain endogenous expression of IL-10 (Maynard et al. 2007). *CD4*^cre−^/^ERT2^-*Tbx21*^flox/++^ (*CD4*^cre−^) and *CD4*^cre+^/^ERT2^-*Tbx21*^flox/++^ (*CD4*^cre+^) mice were generated by crossing *Rosa26*^flSTOPtdRFP^ mice (Luche et al. 2007) with *Tbx21*^flox^ mice (Intlekofer et al. 2008). *Arg1*^flox/++^ and *LysM*^cre^ mice were obtained from The Jackson Laboratory. These mice were bred to generate *LysM*^cre−^*Arg1*^flox/++^ and *LysM*^cre+^*Arg1*^flox/++^ mice. *Arg1*^flox/++^ mice were further bred with *CD4*^cre^ mice to generate *CD4*^cre−^*Arg1*^flox/++^ and *CD4*^cre+^*Arg1*^flox/++^ mice. C57BL/6 WT mice were purchased from Charles River or Envigo.

### Infections and treatments

MCMV was prepared via sorbital gradient purification as described previously (Stacey et al. 2014). Mice were infected with 3 × 10^4^ pfu of MCMV intraperitoneally. *CD4*^cre−^ and *CD4*^cre+^ mice were injected intraperitoneally with tamoxifen (Sigma-Aldrich) as indicated for 5 d from day 7 p.i. at a daily dose of 75 mg/kg (20 mg/ml).

### Next-generation sequencing

Leukocytes were isolated directly *ex vivo* from the SGs of 10BiT mice on day 14 p.i. (minimum *n* = 5 mice per group with three replicates). Pooled cells were labeled using a Zombie Aqua Fixable Viability Kit (BioLegend) and stained with anti-CD16/CD32 (Fc block, BioLegend), anti-CD4–BV605 (clone RM4-5, BioLegend), anti-CD44–FITC (clone IM7, BioLegend), anti-CD62L–PE-Cy7 (clone MEL-14, BioLegend), and anti-CD90/90.1–PE (clone OX-7, BioLegend). Cells were sorted as CD4^+^ CD44^+^ CD62L^−^ CD90/90.1^+^ (Thy1.1^+^) or CD90/90.1^−^ (Thy1.1^−^) populations directly into Buffer RLT or Buffer RLT Plus (Qiagen) using a modified FACS Aria II (BD Biosciences). Total RNA was extracted using an RNeasy Micro Kit or an RNeasy Mini Kit (Qiagen), and RNA integrity scores were determined using an RNA 6000 Pico Kit (Agilent).

### RNA-seq

Multiplexed mRNA libraries were obtained using a TruSeq RNA Library Prep Kit v2 (Illumina) and analyzed using a Bioanalyzer High Sensitivity DNA Chip (Agilent). Libraries were sequenced using a HiSeq 2500 System (Illumina). Paired-end reads (100 bp) were trimmed using Trim Galore (https://www.bioinformatics.babraham.ac.uk/projects/trim_galore/) and assessed for quality using FastQC (https://www.bioinformatics.babraham.ac.uk/projects/fastqc/). Reads were mapped to the mouse GRCm38 reference genome using STAR (Dobin et al. 2013). Counts were assigned to transcripts using featureCounts (Liao, Smyth, and Shi 2014) with the GRCm38.84 Ensembl gene build GTF (http://www.ensembl.org/info/data/ftp/index.html/). Differential gene expression analyses were performed using DESeq2 (Love, Huber, and Anders 2014). Genes were discarded from the analysis if differential expression failed to reach significance (adjusted *P* < 0.05 with Benjamini-Hochberg correction).

### ATAC-seq

ATAC-seq was performed as described previously (Buenrostro et al. 2013) using a Nextera DNA Sample Preparation Kit (Illumina). DNA was isolated after library preparation using a MiniElute PCR Purification Kit (Qiagen). Size selection was performed using a BluePippin System (Sage Science) with 2% Agarose Gel Cassettes (Sage Science). Libraries were sequenced using a HiSeq 4000 System (Illumina). Paired-end reads (100 bp) were trimmed using Trim Galore (https://www.bioinformatics.babraham.ac.uk/projects/trim_galore/) and assessed for quality using FastQC (https://www.bioinformatics.babraham.ac.uk/projects/fastqc/). Reads were mapped to the mouse GRCm38 reference genome using BWA (Li and Durbin 2009). Duplicate reads were removed using MarkDuplicates (Picard) (http://broadinstitute.github.io/picard/). Peaks were called using MACS2 (Zhang et al. 2008) in BAMPE mode (adjusted *P* < 0.05 with Benjamini-Hochberg correction). Differential analysis of open regions was performed using DiffBind (http://bioconductor.org/packages/release/bioc/vignettes/DiffBind/inst/doc/DiffBind.pdf).

### TCR-seq

Unique molecular identifier (UMI)-labeled 5’-RACE TCR cDNA libraries were synthesized synthesis using a Mouse TCR Profiling Kit (MiLaboratories). Indexed samples were pooled and sequenced using a MiSeq System (Illumina). Paired-end reads (150 bp) were extracted and clustered by UMI using MiGEC (Shugay et al. 2014). Sequences were discarded from the analysis if the read count was <4 per cDNA. Error correction was performed using MiGEC (Shugay et al. 2014). Repertoires were extracted using MiXCR (Bolotin et al. 2015). The weighted average use of bulky, charged, and strongly interacting (aromatic and hydrophobic) amino acids positioned centrally in the CDR3β sequences and *TRBV* gene use (weighted by frequency) were calculated using VDJtools (Shugay et al. 2015). Diversity was calculated by downsampling the repertoires to an equal number of UMIs (*n* = 4,300 for TCRα and *n* = 4,000 for TCRβ) three separate times and plotting the mean Chao1 index, reflecting lower bound total diversity, and (1 – normalized Shannon-Wiener index), reflecting clonality. The mean number of unique nucleotide sequences encoding the same amino acid sequence (convergence) was calculated for the 2,000 most prevalent clonotypes in each sample using VDJtools. Overlap was calculated for the 2,000 most prevalent clonotypes using F2 metrics to estimate sharing at the amino acid level among *V* gene-matched sequences in each sample. Cluster analysis was performed using the 500 most prevalent clonotypes in pooled samples (Thy1.1^+^ versus Thy1.1^−^). These datasets were analysed using tcrgrapher (https://github.com/KseniaMIPT/tcrgrapher), an R library based on ALICE (Pogorelyy et al. 2019). The real and expected numbers of neighbors were calculated for each clonotype with a maximum of *n* = 1 amino acid mismatch. Clonotypes with a significantly higher number of neighbors than expected on statistical grounds (adjusted *P* < 0.0001 with Benjamini-Hochberg correction) were identified as tcrgrapher hits. The expected number of neighbors was estimated via generation probabilities calculated for each clonotype using OLGA (Sethna et al. 2019), with the selection factor set at Q = 6.27 (Elhanati et al. 2018).

### Bioinformatics

RNA-seq analysis was performed using DESeq2/1.32.0, dplyr/1.0.7, and GenomicRanges/1.44.0. Volcano plots were drawn using ggplot2/3.3.5 and ggrepel/0.9.1. TCR-seq analysis was performed using MiGEC/1.2.9, MiXCR/3.0.13, VDJtools/1.2.1, tidyverse/1.3.1, igraph/1.2.6, ggnetwork/0.5.10, msa/1.22.0, tcrgrapher/0.0.09000, stringdist/0.9.6.3, and ggseqlogo/0.1. Other software packages used across next-generation sequencing platforms included Nextflow (https://www.nextflow.io/) (Di Tommaso et al. 2017), trimgalore/0.6.4 (https://www.bioinformatics.babraham.ac.uk/projects/trim_galore/), FastQC/0.11.8 (https://www.bioinformatics.babraham.ac.uk/projects/fastqc/), multiqc/1.7 (https://multiqc.info/) (Ewels et al. 2016), STAR/2.7.3 (https://github.com/alexdobin/STAR) (Dobin et al. 2013), BWA/0.7.10 (http://bio-bwa.sourceforge.net/) (Li and Durbin 2009), picard/2.20.2 (http://broadinstitute.github.io/picard/), samtools/1.9 (http://www.htslib.org/) (Danecek et al. 2021), bamtools/2.5.1 (https://github.com/pezmaster31/bamtools) (Barnett et al. 2011), featurecounts/2.0.0 (http://subread.sourceforge.net/) (Liao, Smyth, and Shi 2014), and MACS2/2.1.2 (https://pypi.org/project/MACS2/) (Zhang et al. 2008). Venn diagrams were drawn using InteractiVenn (http://www.interactivenn.net) (Heberle et al. 2015). Heatmaps were drawn using Morpheus (https://software.broadinstitute.org/morpheus). ATACseq data were visualized using Integrative Genomics Viewer (Robinson et al. 2011). Gene ontology analysis was performed using GOTermFinder (https://go.princeton.edu/cgi-bin/GOTermFinder). ATAC-seq motif analysis was performed using The MEME Suite (https://meme-suite.org/meme/doc/cite.html?man_type=web) (Bailey et al. 2015). *Tbx21* binding motifs were obtained using JASPAR^2020^ (Fornes et al. 2020).

### Quantification of MCMV

Infectious virus was quantified in organs via plaque assay as described previously (Stack et al. 2015). Viral loads in oral lavage were quantified via qPCR (Clement et al. 2016; Kamimura and Lanier 2014).

### Western blotting

Leukocytes were isolated as described previously (Stacey et al. 2011). CD4^+^ T cells were purified from SGs and spleens via magnetic separation using a MagniSort Mouse CD4 Positive Selection Kit (Thermo Fisher Scientific). Cell lysates were then generated from equal numbers of cells using NuPAGE™ LDS Sample Buffer (Thermo Fisher Scientific) supplemented with 100 mM dithiothreitol (Sigma-Aldrich). Samples were loaded onto 4–12% NuPAGE Bis-Tris Gels (Thermo Fisher Scientific) after boiling and transferred to a PVDF membrane using an XCell II Blot Module (Thermo Fisher Scientific). Blots were probed with anti-arginase-1 (rabbit polyclonal, Thermo Fisher Scientific) and developed using anti-rabbit IgG–HRP (Bio-Rad). Band intensity was determined using a G:BOX Gel Imaging System (Syngene). Blots were then stripped using Restore PLUS Western Blot Stripping Buffer (Thermo Fisher Scientific) and probed again with anti-actin (rabbit polyclonal, Abcam).

### Flow cytometry

Leukocytes were isolated from SGs and spleens as described previously (Stacey et al. 2011). Cells were labeled using a Zombie Aqua Fixable Viability Kit (BioLegend) and surface-stained with anti-CD16/CD32 (Fc block, BioLegend) and combinations of anti-CD4–BV605, anti-CD4– Pacific Blue, or anti-CD4–APC-Cy7 (clone RM4-5, Biolegend), anti-CD39–Alexa Fluor 647 (clone Duha59, BioLegend), anti-CD44–FITC (clone IM7, BioLegend), anti-CD62L–BV711 or anti-CD62L–PE-Cy7 (clone MEL-14, BioLegend), anti-CD90/90.1–APC, anti-CD90/90.1– BV605, or anti-CD90/90.1–PE (clone OX-7, BioLegend), anti-CD127–BV711 (clone A7R34, BioLegend), anti-PD-1–BV421 (clone 29F.1A12, BioLegend), and anti-Tim-3–BV711 or anti-Tim-3–PerCP-Cy5.5 (clone RMT3-23, BioLegend). Tetramer staining was performed as described previously (Clement et al. 2016). The following specificities were used in this study: H-2D^b^ M45 residues 985–993 (HGIRNASFI), H-2K^b^ IE3 residues 416–423 (RALEYKNL), H-2K^b^ M38 residues 316–323 (SSPPMFRV), and H-2K^b^ m139 residues 419–426 (TVYGFCLL) (National Institutes of Health Tetramer Core Facility). Internal antigen expression was determined after fixation/permeabilization in BD Cytofix/Cytoperm Solution (BD Biosciences) or FoxP3 / Transcription Factor Staining Buffer (eBioscience). Cells were stained with combinations of anti-arginase-1–APC (clone Met1-Lys322, Bio-Techne) or anti-arginase-1– PE-Cy7 (clone A1ex5, eBioscience), anti-Ki-67–Alexa Fluor 488 (clone 11F6, BioLegend), anti-T-bet–BV605 or anti-T-bet–Pacific Blue (clone 4B10, BioLegend), and anti-TCF1/7–Alexa Fluor 647 (clone C63D9, Cell Signaling Technology). Cells were acquired using an Attune NxT (Thermo Fisher Scientific) or an LSR Fortessa (BD Biosciences). All flow cytometry data were analyzed using FlowJo v10.5.3 and v10.8.1 (FlowJo LLC).

### Quantification of anti-SSA

Plasma samples were obtained from cardiac punctures and assessed for anti-Sjögrens syndrome antigen (SSA)/Ro 60 IgG levels via ELISA (Alpha Diagnostics).

### Functional assays

Leukocytes from SGs and spleens were stimulated with peptides at a final concentration of 3 µg/ml for 2 h at 37 °C. The following peptides were used to stimulate CD4^+^ T cells: m09 residues 133–147 (GYLYIYPSAGNSFDL), M25 residues 409–423 (NHLYETPISATAMVI), m139 residues 560–574 (TRPYRYPRVCDASLS), and m142 residues 24–38 (RSRYLTAAAVTAVLQ). The cultures were then supplemented with brefeldin A (2 µg/ml, Sigma-Aldrich) and incubated for a further 4 h at 37 °C. After stimulation, cells were labeled using a Zombie Aqua Fixable Viability Kit (BioLegend) and stained with anti-CD16/CD32 (Fc block, BioLegend) and anti-CD4–APC-Cy7 or anti-CD4–BV605 (clone RM4-5, BioLegend). Internal antigen expression was determined after fixation/permeabilization in Cytofix/Cytoperm Solution (BD Biosciences). Cells were stained with combinations of anti-IFN-γ–APC-Cy7, anti-IFN-γ–FITC, or anti-IFN-γ–Pacific Blue (clone XMG1.2, BioLegend), anti-IL-10–APC or anti-IL-10–FITC (clone JES5-16E3, eBioscience), and anti-T-bet–BV605 (clone 4B10, BioLegend). Cells were acquired using an Attune NxT (Thermo Fisher Scientific), and data were analyzed using FlowJo v10.5.3 (FlowJo LLC).

### Statistics

Sample sizes for next-generation sequencing experiments were calculated using G*Power (https://www.psychologie.hhu.de/arbeitsgruppen/allgemeine-psychologie-und-arbeitspsychologie/gpower.html), where a minimum of *n* = 5 pooled mice per group was used to detect a difference in means with 90% power and an α value set at 0.05 across a minimum of three replicates. All outliers were included in the final datasets. Comparisons between two groups were performed using the Mann-Whitney U test. Significance across all tests is reported as \**P* < 0.05, \*\**P* < 0.01, \*\*\**P* < 0.001, and \*\*\*\**P* < 0.0001.

### Ethics statement

All mouse experiments were approved by the Biological Services Facility (Cardiff University) and performed under UK Home Office Project License P7867DADD.

## AUTHOR CONTRIBUTIONS

M.C., K.L., K.L.M., M.M., L.C., A.C.F., J.S., S.C., and V.V.K. performed experiments; M.C., R.A., V.V.K., K.R.L., O.V.B., D.M.C., D.A.P., and I.R.H., analyzed data; D.R.W. and S.A.J. provided key reagents; I.R.H. directed the study; M.C., O.V.B., D.M.C., D.A.P., and I.R.H wrote the manuscript.

## ACKNOWLEDGEMENTS

We thank Steven Reiner (Columbia University, New York) for providing *Tbx21*^flox^ mice and Joerg Fehling (Ulm University, Germany) for providing *Rosa26*^flSTOPtdRFP^ mice. Biotinylated monomers were obtained from the NIH Tetramer Core Facility. This work was funded by the Medical Research Council via a Confidence in Concept Award to I.R.H., the Ministry of Science and Higher Education of the Russian Federation via a Subsidy Grant to O.V.B. and D.M.C. (075-15-2019-1789), and the Wellcome Trust via Senior Research Fellowships to D.R.W. (110199/Z/15/Z) and I.R.H. (207503/Z/17/Z), a Senior Investigator Award to D.A.P. (100326/Z/12/Z), and a Collaborator Award involving I.R.H. (209213/Z/17/Z).

## EXTENDED DATA FIGURE LEGENDS

**Extended Data Figure 1.**
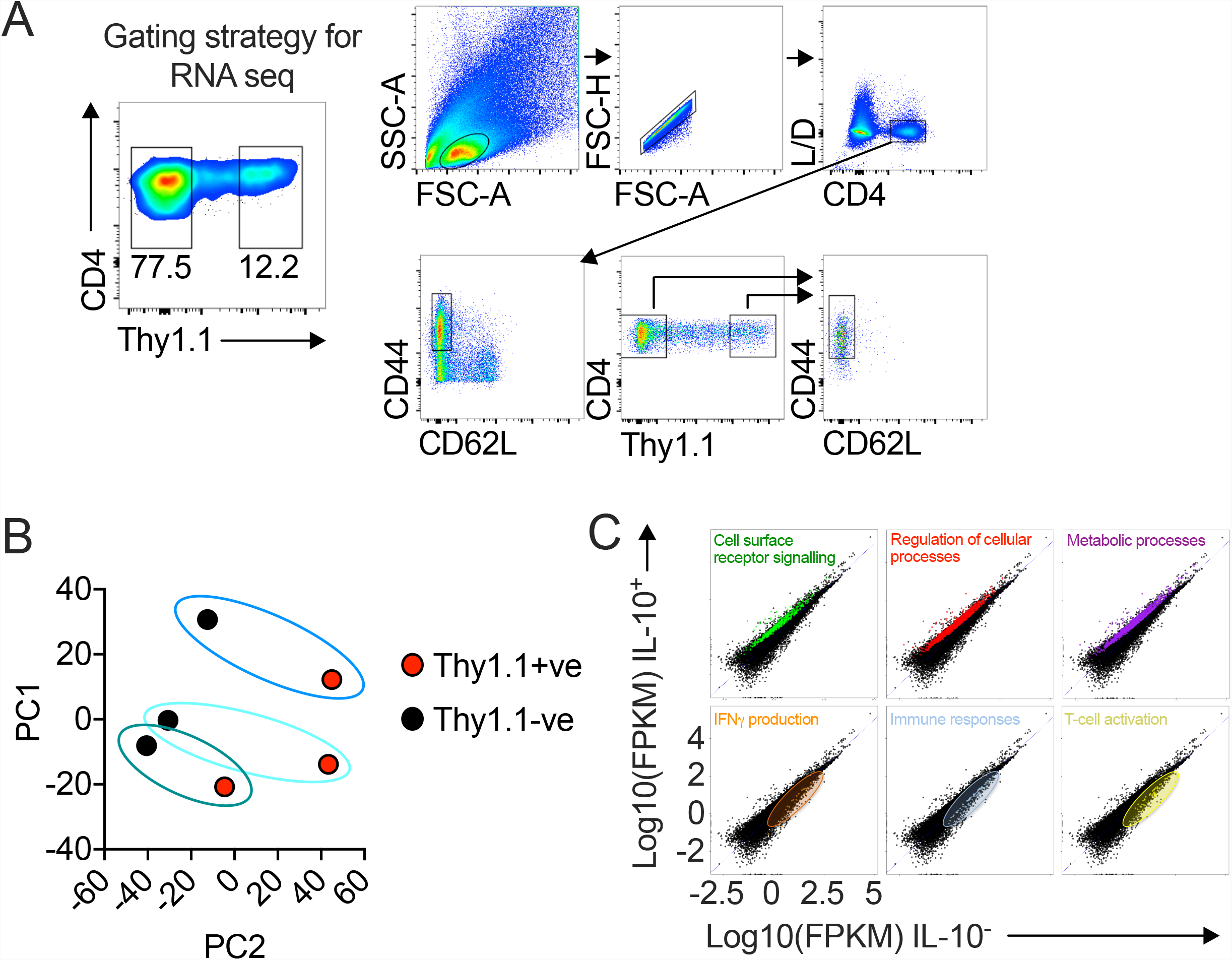
Gating strategy for RNA-seq, ATAC-seq, and TCR-seq analysis of CD4^+^ T cells isolated from the SGs. (**a**) 10BiT mice were infected with 3 × 10^4^ pfu of MCMV. Leukocytes were isolated from the SGs on day 14 p.i. and sorted as CD4^+^ CD44^+^ CD62L^−^ CD90/90.1^+^ (Thy1.1^+^) or CD90/90.1^−^ (Thy1.1^−^) populations via FACS. Data are shown as pooled analyses from a minimum of *n* = 5 mice per group representing three independent experiments. (**b**) PCA plots from (**a**) after RNA-seq analysis. (**c**) Gene expression levels identifying the location of gene clusters from the gene ontology analysis (**Fig. 1c**).

**Extended Data Figure 2.**
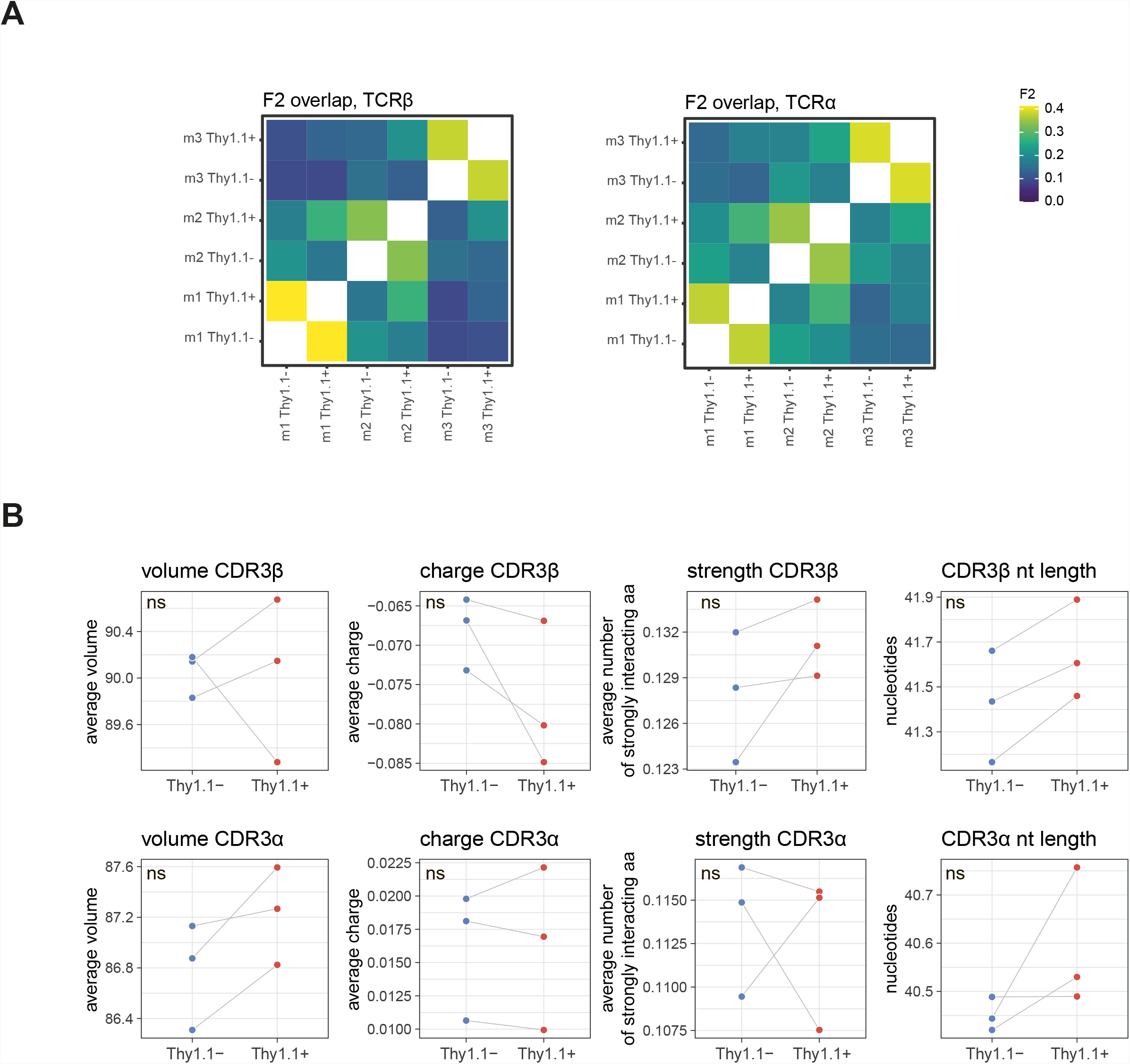
Analysis of repertoire overlap and the physicochemical properties of TCRs. (**a**) F2 overlap metrics for the 2,000 most prevalent clonotypes showing pairwise calculations to estimate the sharing of amino acid sequences with identical *V* genes. Left: TCRβ. Right: TCRα. (**b**) Mean physicochemical properties of the central five amino acids in the CDR3β (top) and CDR3α loops (bottom) weighted by clonotype frequency. Right: averaged and weighted nucleotide lengths for CDR3β (top) and CDR3α (bottom). Significance was evaluated using a paired t-test with Benjamini-Hochberg correction. ns, not significant.

**Extended Data Figure 3.**
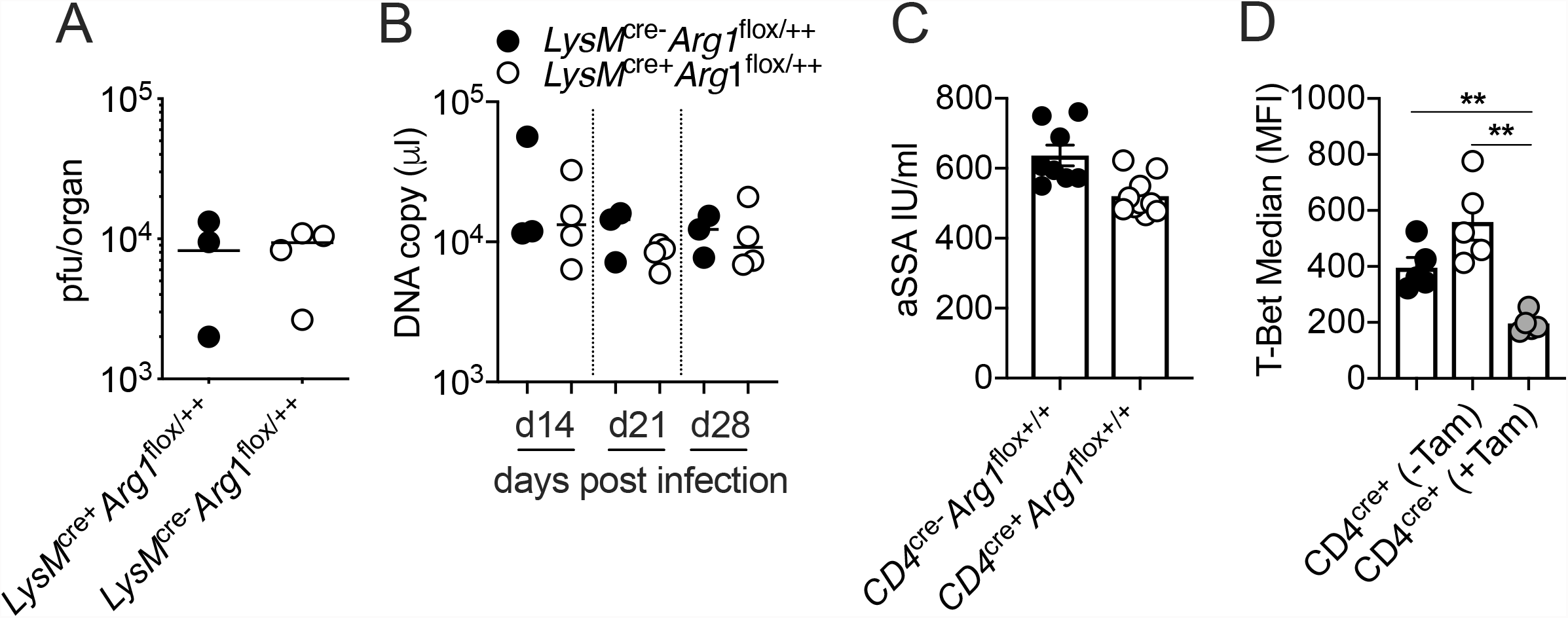
Arg1 expression in myeloid cells does not impact the replication of MCMV. *LysM*^cre−^*Arg1*f^lox+/+^ and *LysM*^cre+^*Arg1*f^lox+/+^ mice were infected with 3 × 10^4^ pfu of MCMV. (**a**) MCMV replication in SG homogenates on day 28 p.i. measured via plaque assay. Data are shown as individual points with median values (*n* = 3–4 mice per group representing two independent experiments). (**b**) Viral genomes in saliva on days 14, 21, and 28 p.i. measured via qPCR. Data are shown as individual points with median values (*n* = 3–4 mice per group representing two independent experiments). (**c**) *CD4*^cre−^*Arg1*^flox+/+^ and *CD4*^cre+^*Arg1*^flox+/+^ mice were infected with 3 × 10^4^ pfu of MCMV. Cardiac punctures were performed on day 30 p.i. Anti-SSA IgG titers were measured in plasma samples via ELISA. Data are shown as mean ± SEM (*n* = 10 mice per group representing two independent experiments). (**d**) *CD4*^cre−^/^ERT2^-*Tbx21*^flox/++^ (*CD4*^cre−^) and *CD4*^cre+^/^ERT2^-*Tbx21*^flox/++^ (*CD4*^cre+^) mice were infected with 3 × 10^4^ pfu of MCMV. Tamoxifen was administered (+Tam) or withheld (−Tam) from days 7 to 12 p.i. Leukocytes were isolated from the SGs on day 14 p.i. The expression of T-bet was measured via flow cytometry. Data are shown as mean ± SEM (*n* = 4–5 mice per group representing four or five independent experiments). MFI, median fluorescence intensity. \*\**P* < 0.01 (Mann-Whitney U test).

